# A gradual backward shift of dopamine responses during associative learning

**DOI:** 10.1101/2020.10.04.325324

**Authors:** Ryunosuke Amo, Akihiro Yamanaka, Kenji F. Tanaka, Naoshige Uchida, Mitsuko Watabe-Uchida

## Abstract

It has been proposed that the activity of dopamine neurons approximates temporal difference (TD) prediction error, a teaching signal developed in reinforcement learning, a field of machine learning. However, whether this similarity holds true during learning remains elusive. In particular, some TD learning models predict that the error signal gradually shifts backward in time from reward delivery to a reward-predictive cue, but previous experiments failed to observe such a gradual shift in dopamine activity. Here we demonstrate conditions in which such a shift can be detected experimentally. These shared dynamics of TD error and dopamine activity narrow the gap between machine learning theory and biological brains, tightening a long-sought link.

## Introduction

Maximizing future reward is one of the most important objectives of learning necessary for survival. To achieve this, animals learn to predict the outcome associated with different objects or environmental cues, and optimize their behavior based on the predictions. Dopamine neurons play an important role in reward-based learning. Dopamine neurons respond to an unexpected reward by phasic excitation. When animals learn to associate a reward with a preceding cue, dopamine neurons gradually decrease responses to the reward itself^1,2^. Because these dynamics resemble the prediction error term in animal learning models such as that in the Rescorla-Wagner model^3^, it is thought that dopamine neurons broadcast reward prediction errors (RPEs), the discrepancy between actual and expected reward value, to support associative learning.

In machine learning, the temporal difference (TD) learning algorithm is one of the most influential algorithms, which uses a specific form of teaching signal, called TD error^4^. The aforementioned Rescorla-Wagner model treats an entire trial as a single discrete event, and does not take into account timing within a trial. In contrast, TD learning considers timing within a trial, and computes moment-by-moment prediction errors based on the difference (or change) in values between consecutive time points in addition to rewards received at each moment. TD learning models explain dopamine neurons’ responses to reward-predictive cues as a TD error; a cue-evoked dopamine response occurs because a reward-predictive cue indicates a sudden *increase* in value.

Despite these remarkable resemblance between the activity of dopamine neurons and TD error signals, a key prediction of TD errors – whether the activity of dopamine neurons follows TD learning models during learning^1,2^ – remains elusive. One of the hallmarks of TD learning algorithms is that the value as well as TD errors gradually shifts backward in time from the reward to earlier points^2,5^. In other words, in standard TD models, learning happens incrementally by gradually shifting a reward-associated value to earlier and earlier time points so, after learning, value can be inferred as soon as predictive information is available. As the value gradually shifts to earlier time points, TD errors also moves backward in time because an increase in value occurs earlier in time. However, this most characteristic signature of TD errors, a temporal gradual shift of signals over learning, has not been observed in dopamine activity, despite previous attempts^6–9^. On one hand, the lack of such a unique signature of TD (even if not all TD errors show gradual shifts^6,10^) together with other observations indicating complexities of dopamine activity has encouraged alternative theories for dopamine^11,12^ which reject a comprehensive account of TD learning. On the other hand, recent findings reinforce the idea that dopamine activity follows TD errors even for non-canonical activity, such as ramping dopamine signals observed in dynamic environments^13^ and the dynamic dopamine signal that tracks moment-by-moment changes in the expected reward (value) that unfold as the animal goes through different mental states within a trial^14^.

There are multiple possible implementations of TD models, some of which predict that TD errors show gradual temporal shifts and some of which do not require this to happen, depending on the learning strategy used^6,15–17^. In other words, TD errors show a gradual temporal shift when an agent takes a specific learning strategy. Thus, depending on the experimental design, we may either observe a gradual temporal shift in dopamine activity, or totally miss it. Here, we employed different task conditions and examined temporal dynamics of dopamine cue responses that allowed us to observe this long-sought gradual shift. We also modelled and explored the conditions necessary for small shifting signals to be enhanced, and thus easily detected.

## Results

### Dynamics of dopamine cue responses for first time learning of cue-reward association

The activity of dopamine neurons during associative learning has typically been studied using well-trained animals, and observations of dopamine activity during the learning phase where the temporal shift is predicted to occur have been limited^1,6,18–20^. We previously recorded population activity of dopamine axons in the ventral striatum (VS), a major target area of dopamine projections, using fiber-fluorometry (photometry) while naive animals learned, for the first time, to associate odor cues and water reward (classical conditioning) in a head-fixed preparation^7^. During learning in naive animals, dopamine axons showed an activity change characteristic of RPE, i.e. increase of cue responses and decrease of reward responses, although we could not observe a clear temporal shift of dopamine activity. This was potentially caused by insufficient temporal resolution due to the slow kinetics of GCaMP6m^21^, the Ca^2+^ indicator used in the previous study^7^. In the present study, we simply increased the delay duration between the cue and the reward (Figure 1). To detect dopamine dynamics, we injected adeno associated virus (AAV) in the VS to express the dopamine sensor GRAB_DA2m_ (DA2m)^22^ and measured fluorescence changes with fiber-fluorometry (Figure 1a, Supplemental Figure 1). Over multiple sessions of classical conditioning, naive mice gradually acquired anticipatory licking (Figure 1b). At the same time, as expected, dopamine responses to the reward-predicting cue gradually developed (Figure 1c), whereas responses to reward gradually decreased (Figure1c, d). In contrast, responses to an unexpected reward (free reward) stayed high throughout a session (Figure 1c, d).

**Figure 1.**
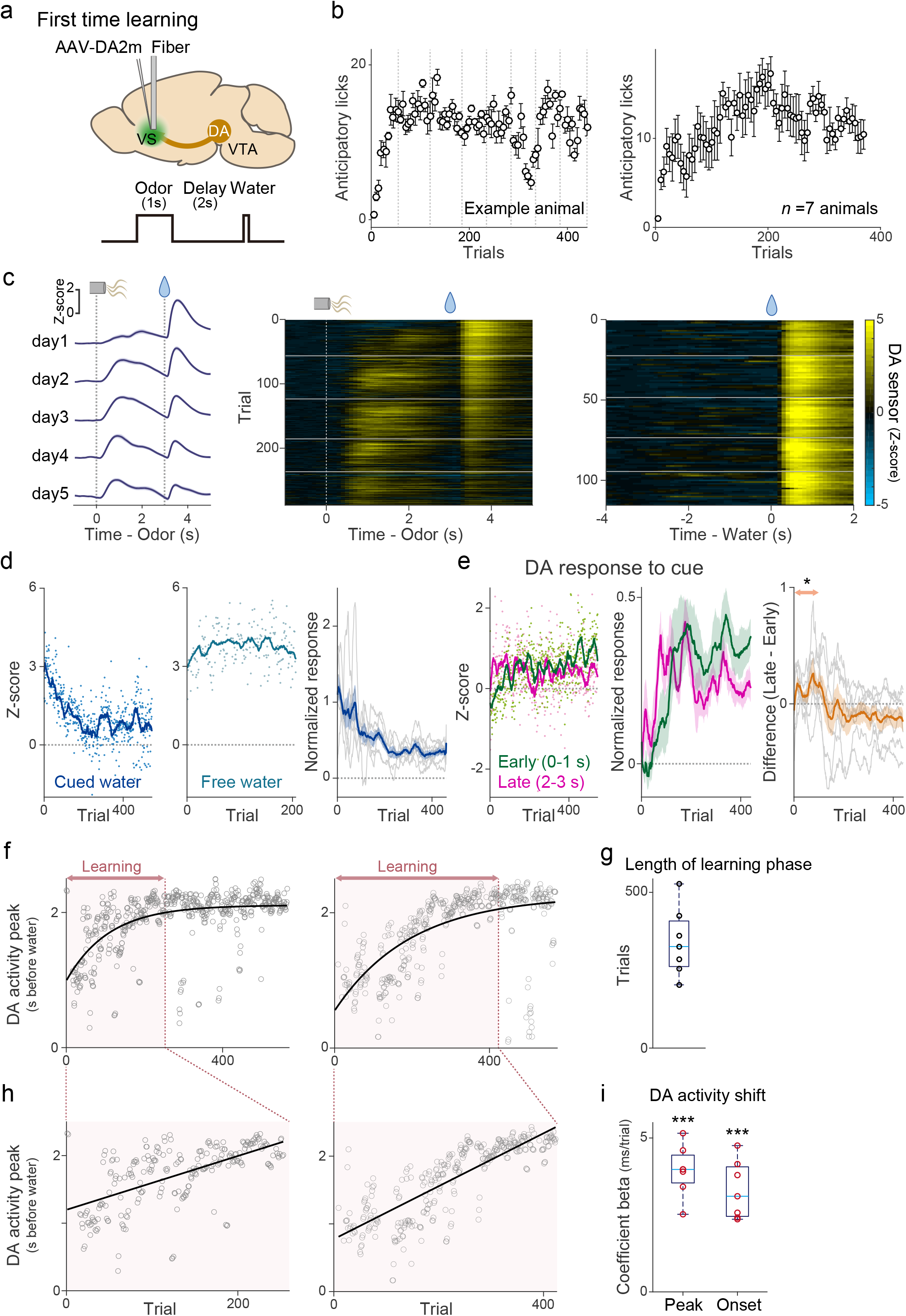
Dopamine release in the ventral striatum during first-time classical conditioning. (a) Both dopamine sensor (DA2m) and optical fiber for fluorometry were targeted to VS. (**b**) Lick counts (5 trials mean ± sem) during the delay period (0-3 s after odor onset). Dotted lines (gray) indicate boundaries of sessions. (**c**) Dopamine signals for cued water trials (left, mean ± sem, and center) and for free water trials (right) in an example mouse. Horizontal lines (white) indicate boundaries of sessions. (**d**) Dopamine responses to cued water (left) and to free water (center) in an example mouse, and responses (normalized with free water responses) to cued water in all animals (right, gray: each animal; blue: mean ± sem). Each dot (left, center) represents responses in each trial, and a line shows moving averages of 20 trials. (**e**) Responses to a reward-predicting odor in an example animal (left) and in all animals (center, mean ± sem). early: 0-1 s from odor onset (green); late: 2-3 s from odor onset (magenta). Right, difference between early and late odor responses (grey: each animal; orange: mean ± sem). Dopamine activity in the early phase was significantly higher than activity in the late phase during the first 2-100 trials (*t* = 3.3, *p* = 0.017). (**f**) Peak timing of dopamine sensor signal during the delay period (gray circles) were fitted with exponential curve (black) in 2 example animals. The learning phase was determined by trials with more than 1ms/trial of the temporal shift of the peak (red). (**g**) Length of the learning phase in all animals. (**h**) Linear regression of peak timing of dopamine sensor signal during learning phase (circles) with trial number. (**i**) Regression coefficients for peak timing and onset timing of dopamine sensor signal with trial number. Regression coefficients for both peak and onset were significantly positive (*t* = 13, *p* = 0.16 × 10^−4^ for peak, *t* = 9.3, *p* = 0.88 × 10^−4^ for onset; t-test). Red circle, significant slopes (*p*-value ≤ 0.05). n=7 animals.

Looking at patterns of dopamine activity in each animal closely, we noticed some dynamical changes (Figure 1c); over learning, activity during the delay period (after cue onset to water delivery) was systematically altered. We observed that dopamine excitation was more prominent in the later phase of the delay period (Figure 1c, e Late) early on in training, whereas after learning cue responses were typically observed immediately following the cue onset (Figure 1c, e Early). Indeed, during early learning, activity during the late phase of the delay period was significantly higher than activity during the early phase of the delay period (Figure 1e).

In order to examine the temporal dynamics of dopamine activity in greater detail, we first tested how the peak of activity during the delay period changed over learning. We plotted the time point when the dopamine signal reached its maximum before water delivery in each trial (Supplemental Figure 2). Next, we fitted an exponential function to the peak timing plotted as a function of trial number because the timing of the peak plateaued after a certain number of trials. In each animal, we observed a consistent shift of the peak timing (Figure 1f). In Figure 1h, we zoomed in on the learning phase, by excluding the later phase where the peak timing had plateaued (did not change any more than 1 ms/trial based on the exponential fit). The length of the learning phase was variable across animals (Figure 1g). However, we found that the temporal shift of the response peak was reliably observed in all animals, with the average shift of about 4 ms/trial during the learning phase (Figure 1h, i). This temporal shift was also detected when we analyzed the timing of the excitation onset instead of the peak (Figure 1i, Supplemental Figure 2b, c). Together, we detected a significant temporal shift of dopamine activity in all the animals we examined both in terms of the peak and the excitation onset before reward.

### Dynamics of cue responses in dopamine axons in reversal tasks

The above experiment showed that there is a gradual temporal shift of dopamine activity while naive mice learned an association between an odor and reward. However, it remains unknown whether a shift occurs in other conditions, in particular, in well-trained animals with which previous experiments did not report a temporal shift^7,17,18^. We therefore next examined dopamine dynamics in well-trained animals. Specifically, we trained mice in a classical conditioning task for more than 12 days, and then examined dopamine axon activity or dopamine concentration in VS while the mice learned a new association (Figure 2). We focused on a reversal task, where a familiar odor that was previously associated with no outcome became associated with reward (Figure 2a), in order to avoid using a novel cue that can cause small excitation in some dopamine neurons^23,24^. The well-trained mice developed anticipatory licking within the first session of reversal (Figure 2b), faster than the aforementioned initial learning (Figure 1b). When we examined dopamine axon activity in each animal, the dopamine signal showed the temporal shift during the delay period (Figure 2c, d), similar to the initial learning but at a faster speed. We quantified the timing of the peak activity on the first day of learning, and found that the peak timing of both dopamine axon activity and dopamine concentration in VS showed a positive correlation with the trial number (Figure 2e, Supplemental Figure 3, 4a). The temporal shift of dopamine axon activity was also observed in the reversal task in which we switched an outcome from aversive air puff to water (Figure 2e). As a population, the peak timing of average dopamine axon activity and average dopamine concentration showed a significant correlation with the trial number during the reversal tasks (Figure 2f, Supplemental Figure 4).

**Figure 2.**
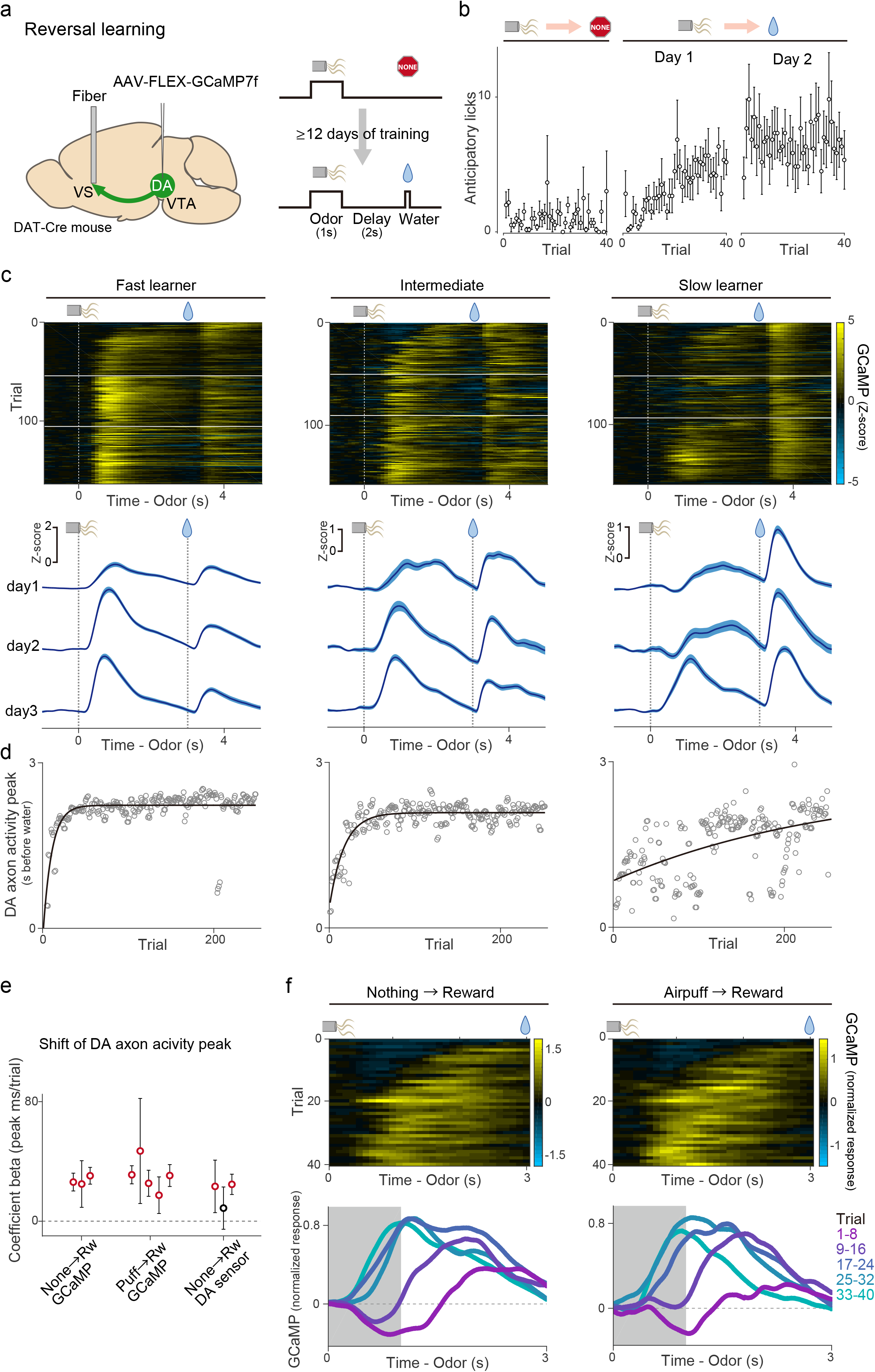
Dopamine axon activity in reversal learning. (**a**) AAV-GCaMP was injected into the VTA, and an optical fiber for fiber-fluorometry was targeted to VS. (**b**) Lick counts during the delay period (0-3 s after odor onset) with reversal training from nothing to reward (n = 6 animals, 3 animals with GCaMP and 3 animals with DA sensor were pooled). (**c**) Dopamine axon activity in 3 sessions after reversal from nothing to reward in 3 example animals. Horizontal white lines (top) indicate boundaries of sessions. Bottom, mean ± sem. (**d**) Dopamine axon activity peak (gray circles) and exponential curve fitted to peak across trials (black). (**e**) Regression coefficient ±95% confidence intervals between activity peak and trial number in each animal in different experimental conditions. Red circles, significant (*p* ≤ 0.05) slopes. (**f**) Dopamine axon activity (normalized with free water response) in the first day of reversal from nothing to reward (left, n = 3 animals) and from airpuff to reward (n = 5 animals). Bottom, each line shows 8 trials mean population neural activity across the session. Gray patches indicate odor presenting periods.

Signals recorded using fiber-fluorometry are inevitably contaminated by artifacts caused by movement. To exclude possible contributions of motion artifacts in our results, we examined the relationship between dopamine signals and licking behavior, which is a major source of movement artifacts in recording in head-fixed animals (Supplemental Figure 5). We first examined the peak timing of anticipatory licking in each trial. The results showed that anticipatory licks during learning peaked *later* than dopamine activity peaks (877 ± 180 ms slower than dopamine activity peak). Next, we analyzed the timing of the initiation of anticipatory licking. The first lick also appeared later than dopamine responses in many trials (427 ± 241 ms slower than dopamine activity peak, and 989 ± 154 ms slower than dopamine excitation onset) (Supplemental Figure 5e, g). With learning, animals tended to increase the vigor of anticipatory licking (total number of licks per trial) (Supplemental Figure 5b, c). On the other hand, we did not observe consistent temporal changes of the lick timing (Supplemental Figure 5d, f). As a result, we did not see a trial-to-trial correlation between the timing of the dopamine activity peak and the timing of initiation or peak of anticipatory licking (Supplemental Figure 5e, h). Thus, licking behaviors cannot explain the observed dynamics of dopamine signals in our recording. To control for any other potential recording artifacts, we also measured fluctuation of signals of the control fluorescence (red fluorescent protein, tdTomato) simultaneously with recording of GCaMP signals in some animals (Supplemental Figure 5a, f). tdTomato signals showed a different pattern of fluctuation compared to the GCaMP signals, and were not consistent across animals, suggesting that the recording artifacts cannot explain the dopamine activity pattern observed in this experiment. Importantly, while a significant temporal shift was observed in the dopamine activity peak, neither tdTomato signal peaks nor the peaks or initiation of anticipatory licks showed a significant shift (Supplemental Figure 5f).

### Dopamine dynamics in repeated associative learning

Our observation indicates that not only initial learning in naïve animals, but also reward learning with a familiar odor in well-trained animals exhibit a backward temporal shift of dopamine activity. What about learning from a novel cue, which is typically used in most experiments^18,25^? We examined dopamine axon activity during learning of a novel odor-water association in well-trained animals (Figure 3). As we mentioned above, multiple studies found that some dopamine neurons respond to a novel stimulus with excitation, especially when using a cue with the same modality as a previously learned rewarding cue, likely due to generalization^23,26,27^. We also observed small excitation to a novel odor in dopamine axon activity in well-trained animals, which presumably corresponds to generalization of initial value (Figure 3a, Supplemental Figure 6). In addition to the transient excitation at the onset of a cue, we found that there was another excitation during the later phase of the delay period (Figure 3a). If we focused only on the later phase of dopamine activity, the excitation appeared to be shifting backward in time. Hence, we quantified the peak timing. Because there were two peaks in some trials, in this case, we detected up to two peaks instead of detecting only the maximum (see Methods) (Figure 3b). When we examined the timing of the later peak (the first peak in case of one peak or the second peak in case of two peaks) during the delay period, we observed a significant correlation between the peak timing and the trial number (Figure 3c), indicating a temporal shift over time.

**Figure 3.**
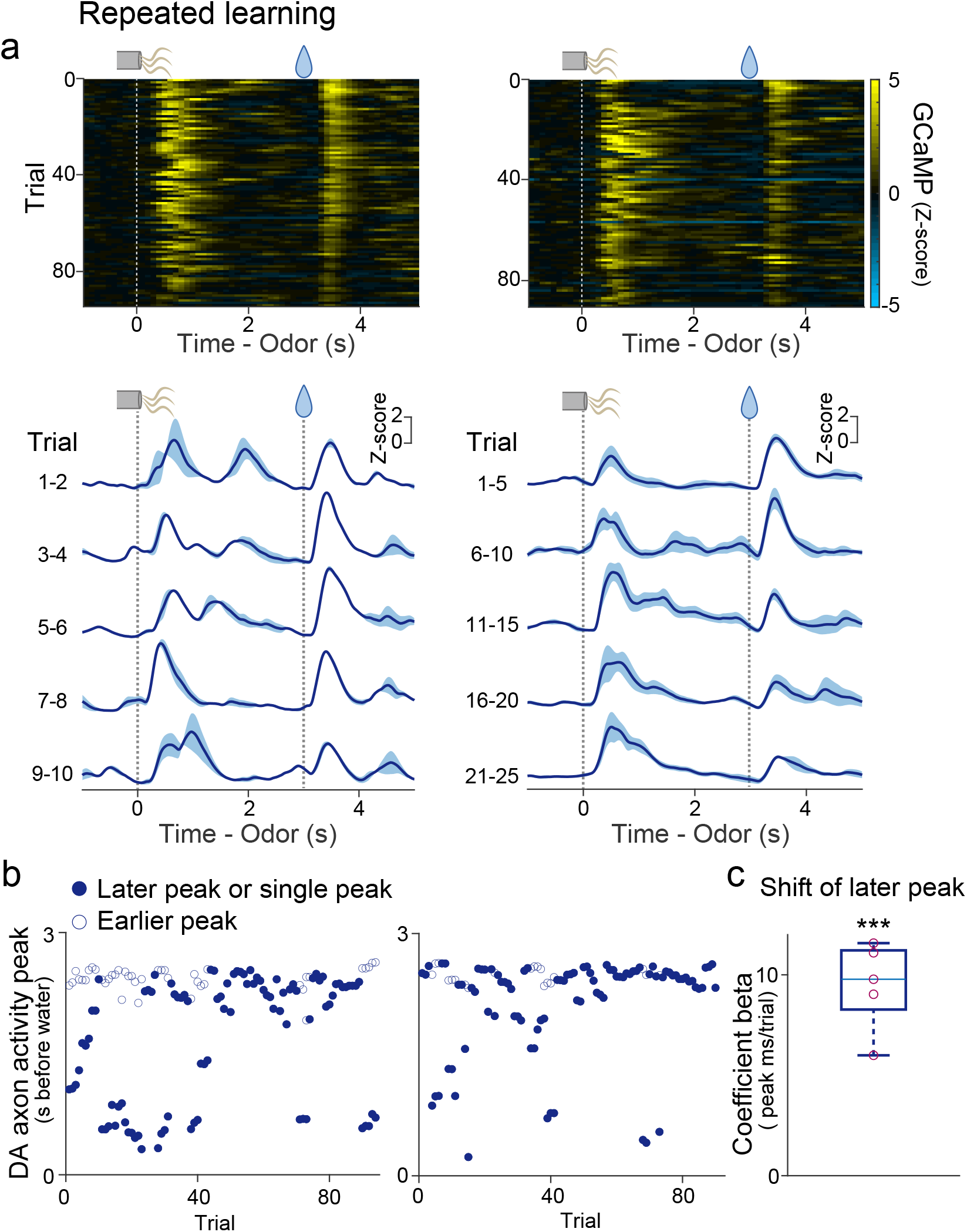
Dopamine axon activity in repeated associative learning. (**a**) GCaMP activity in odor-reward association trials in 2 example animals. Bottom, mean ± sem. (**b**) GCaMP activity peaks (up to 2 for each trial) in the same animals. Filled circles represent the 2nd peak (or peak in trials with only one). (**c**) Linear regression coefficients for 2nd peak timing with trial number (n = 5 animals; *t* = 9.6, *p* = 0.65 × 10^−3^; t-test). Red circles, significant (*p*-value ≤ 0.05, *F*-test).

## Discussion

Our results show that a temporal shift of dopamine activity can be detected in various simple learning paradigms, although we found that it was important to examine in each animal closely because of the variability of shifting speeds and other confounding factors. For example, it can be more difficult to detect a shift when there is an additional factor that may affect dopamine activity such as generalization^26,27^ (Figure 3). Of note, in most cases, dopamine responses to water decreased gradually over learning, but did not totally disappear, similar to many previous studies, likely due to the temporal uncertainty because of the delay between cue and reward^28^.

Although many forms of TD models predict a temporal shift of the TD error signal, previous studies have failed to detect such a shift in dopamine activity during learning. What could prevent the detection of a temporal shift? First, we consider various differences in recording methods. Fluorometry signals using Ca^2+^ indicator and dopamine sensor show slower kinetics than spike data, and can be approximated by a slowly convolved version of the underlying spiking activity. Our simulation analysis indicated that the mere convolution of TD errors with a filter mimicking fluorometry signals exaggerates small signals during the temporal shift (Figure 4). Of note, the temporal shift is not observed by convolution when the original model does not exhibit a temporal shift, such as a learning model involving a Monte-Carlo update (Figure 4a, b). In contrast, TD model (TD-λ in this example; see Methods) show a very small “bump” shifting backward in time in the original model (Figure 4c, d). Because these bumps, although small, spread over some time windows, a convolution with a slow filter (kernel)^13^ accumulates these signals over time, and exaggerates them (Figure 4c), resulting in more “visible” bumps exhibiting a temporal shift (Figure 4d). The use of a slow measurement such as fluorometry can, thus, facilitate the detection of a temporal shift even when the amplitude of each bump is very small. To compensate for the slowness, it was also necessary for us to increase the delay between cue and reward compared to previous studies^7^. Additionally, fluorometry records population activity from a large number of neurons of a specific cell-type, which could make detection of common activity features across neurons easier by averaging. These two factors, slow signals and averaging across the population, can increase the likelihood of detecting a temporal shift. Future studies with single cell resolution will address how TD error signals manifest at a single neuron level.

**Figure 4.**
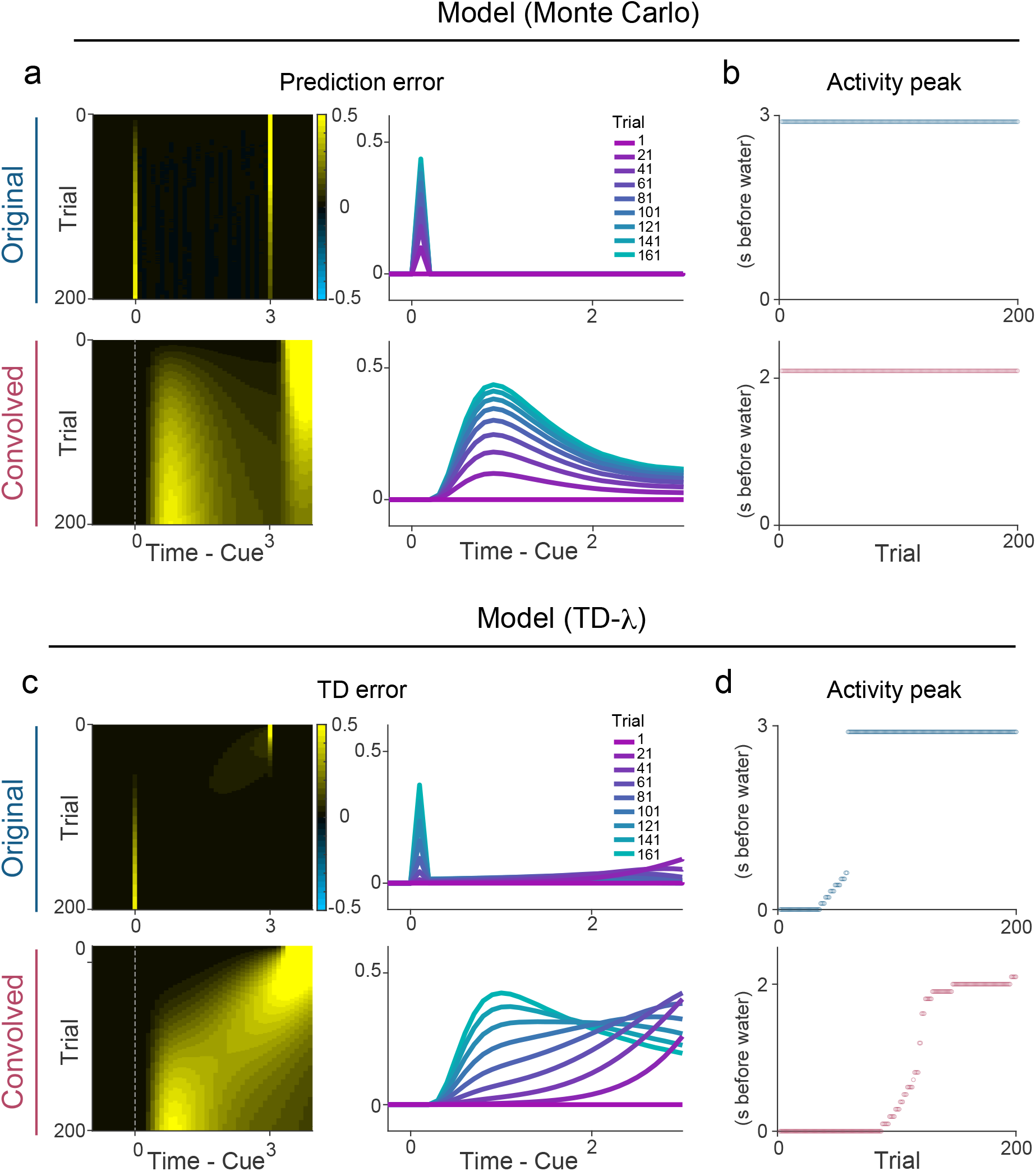
Dynamics of prediction error signals in models with different update rules. (**a**) Prediction error signals during odor-reward associative learning with Monte Carlo updates (top) and signals convolved with GCaMP-like kernel to mimic fluorometry signals (bottom). (**b**) Peak timing of prediction error signals during a delay period. (**c**) Prediction error signals in TD-λ model (top) and signals convolved to mimic fluorometry signals (bottom). (**d**) Peak timing of prediction error signals during a delay period. Convolution exaggerates a peak shift.

Second, it is important to consider both heterogeneity of dopamine neurons and training history. In the present study, we focused on the dopamine axon activity/concentration in VS, because the dopamine axon activity in VS tends to show canonical RPE signals^7^. Detecting a shift in a novel odor-reward association in well-trained animals was still not straightforward. If animals are trained further, their strategy of learning can improve by changing a parameter in learning such as “λλ” (lambda, a parameter for attention or eligibility trace^6^ in TD models), by suppressing behaviorally irrelevant distinctions between states (state abstraction^15,16^) during delay periods, and/or by inferring states from past learning experiences (belief state^17,29^). In such a case of over-training, the predicted value may not slowly shift but shift immediately to the cue onset. Accordingly, dopamine neurons in animals with different training histories may show a varying extent of temporal shift. In addition, subpopulations of dopamine neurons with different projection targets potentially use a slightly different learning strategy, which would result in different levels of temporal shift. In the future, detection of a temporal shift in dopamine activity may be used to estimate a learning strategy of animals, and of brain areas.

The incremental temporal shift of value is the hallmark of TD learning algorithm which provides a solution to credit assignment problems through bootstrapping of value updates^30^. Despite the powerfulness of TD learning algorithms in machine learning^31,32^, the signature of the temporal shift had not been observed in the brain. This has often been taken as evidence against dopamine activity as a TD error signal, and has hampered our understanding of how dopamine regulates learning in the brain. Here, we observed the actual temporal shift of dopamine responses during learning both in naïve and well-trained animals. These observations provide a foundation for understanding how dopamine functions in the brain, and ultimately deeper understanding of computational algorithms underlying reinforcement learning in the brain.

## Method

### Animal

12 both female and male mice, 2-7 months old were used in this study. We used heterozygote for DAT-Cre (Slc6a3^tm1.1(cre)Bkmn^; The Jackson Laboratory, 006660)^33^, DAT-tTa (Tg(Slc6a3-tTA)2Kftnk; this study), VGluT3-Cre (Slc17a8^tm1.1(cre)Hze^; The Jackson Laboratory, 028534)^34^, and LSL-tdTomato (Gt(ROSA)26Sor^tm14(CAG-tdTomato)Hze^; The Jackson Laboratory, 007914)^35^ transgenic lines. VGluT3-Cre lines crossed with LSL-tdTomato were used for experiments with DA sensor, without use of Cre recombinase. Mice are housed on a 12 hr dark (7:00-19:00)/ 12 hr light (19:00-7:00) cycle. Experiments were performed in the dark period. All procedures were performed in accordance with the National Institutes of Health Guide for the Care and Use of Laboratory Animals and approved by the Harvard Animal Care and Use Committee

### Generation of Slc6a3-tTA BAC transgenic mice

Mouse BAC DNA (clone RP24-158J12, containing the Slc6a3 gene, also known as dopamine transporter gene) was modified by BAC recombination. A cassette containing the mammalianized tTA-SV40 polyadenylation signal^36^ was inserted into the translation initiation site of the Slc6a3 gene. The modified BAC DNA was linearized by PI-SceI enzyme digestion (NEB, USA) and injected into fertilized eggs from CBA/C57BL6 mice. We obtained 3 founders (lines 2, 10 and 15) and selected line 2 for the Slc6a3-tTA mouse line due to higher transgene penetrance in the dopamine neurons.

### Plasmid and virus

To make pAAV-TRE3G-GCaMP6f-WPRE, we first made pAAV-TRE3G-WPRE by inserting WPRE from pAAV-EF1a-DIO-hChR2(H134R)-EYFP-WPRE^37^ (gift from Karl Deisseroth; addgene, #20298) cleaved with ClaI and blunted, into pAAV-TRE3G-GCaMP6^38^, cleaved with EcoRI and BglII and blunted to remove GCaMP6 and Flex site. Then GCaMP6f from pGP-CMV-GCaMP6f (gift from Douglas Kim & GENIE Project; addgene, #40755)^39^ cleaved with BglII and NotI and blunted is inserted into pAAV-TRE3G-WPRE cleaved with ApaI and SalI and blunted to remove extra loxP site. Plasmids of pAAV-TRE3G-GCaMP6f-WPRE and pGP-AAV-CAG-FLEX-jGCaMP7f-WPRE (gift from Douglas Kim & GENIE Project; addgene, #104496)^40^ were amplified and purified with endofree preparation kit (Qiagen) and packaged into AAV at UNC vector core.

### Surgery for virus injection, head-plate installation, and fiber implantation

The surgery was performed under aseptic conditions. Mice are anesthetized with isoflurane (1-2% at 0.5-1 L/min) and local anesthetic (lidocaine (2%)/bupivacaine (0.5%) 1:1 mixture, S.C.) was applied at the incision site. Analgesia (ketoprofen for post-operative treatment, 5 mg/kg, I.P.; buprenorphine for pre-operative treatment, 0.1 mg/kg, I.P.) was administered for 3 days following surgery. Custom-made head-plate was placed on the well-cleaned and dried skull with adhesive cement (C&B Metabond, Parkell) containing a small amount of charcoal powder. To express GCaMP in the dopamine neurons, AAV5-CAG-FLEX-GCaMP7f (1.8 × 10^13^ particles/ml) and AAV5-TRE3G-GCaMP6f (8 × 10^12^ particles/ml) were injected unilaterally in the VTA (500 nl, Bregma -3.1 mm AP, 0.5 mm ML, 4.35 mm DV) in 3 DAT-Cre mice and 2 DAT-tTA mice, respectively. In experiments of reversal learning and repeated learning, AAV5-CAG-FLEX-tdTomato (7.8 × 10^12^ particles/ml; UNC Vector Core) and AAV5-CAG-FLEX-GCaMP7f were co-injected (1:1) in 3 DAT-cre mice and, AAV5-CAG-tdTomato (4.3 × 10^12^ particles/ml; UNC Vector Core) and AAV5-TRE3G-GCaMP6f were co-injected (1:1) in 2 DAT-tTA mice. For expression of dopamine sensor in the VS, AAV9-hSyn-DA2m (1.01 × 10^13^; ViGene bioscience)^22^ was injected unilaterally in the VS (300nl, Bregma +1.45 AP, 1.4 ML, 4.35 DV) in 7 VGluT3-Cre/LDL-tdTomato mice. These mice with dopamine sensor were used for experiments of initial learning, and 3 of them were also used for reversal learning. A glass pipette containing AAV was slowly moved down to the target over the course of a few minutes and kept for 2 minutes to make it stable. AAV solution was slowly injected (∼15 min) and the pipette was left for 10-20 min. Then the pipette was slowly removed over the course of several minutes to prevent the leak of virus and damage to the tissue. An optical fiber (400 μm core diameter, 0.48 NA; Doric) was implanted in the VS (Bregma +1.45 mm 1.4 mm from the midline, 4.15 mm deep from dura). The fiber was slowly lowered to the target and fixed with adhesive cement (C&B Metabond, Parkell) containing charcoal powder to prevent contamination of environmental light and leak of laser light. A small amount of rapid-curing epoxy (Devcon, A00254) was applied on the cement to glue the fiber better.

### Fiber-fluorometry (photometry)

To effectively collect the fluorescence signal from the deep brain structure, we used custom-made fiber-fluorometry as described^41^. Blue light from 473 nm DPSS laser (Opto Engine LLC) and green light from 561 nm DPSS laser (Opto Engine LLC) were attenuated through neutral density filter (4.0 optical density, Thorlab) and coupled into an optical fiber patch cord (400 μm, Doric) using 0.65 NA 20x objective lens (Olympus). This patch cord was connected to the implanted fiber to deliver excitation light to the brain and collect the fluorescence emission signals from the brain simultaneously. The green and red fluorescence signals from the brain were spectrally separated from the excitation lights using a dichroic mirror (FF01-493/574-Di01, Semrock). The fluorescence signals were separated into green and red signals using another dichroic mirror (T556lpxr,Chroma), passed through a band pass filter (ET500/40x for green, Chroma; FF01-661/20 for red, Semrock), focused onto a photodetector (FDS10×10, Thorlab), and connected to a current amplifier (SR570, Stanford Research systems). The preamplifier outputs (voltage signals) were digitized through a NIDAQ board (PCI-e6321, National Instruments) and stored in computer using custom software written in LabVIEW (National Instruments). Light intensity at the tip of patchcord was adjusted to 200 μW and 50 μW for GCaMP and DA2m, respectively.

### Histology

All mice used in the experiments were examined for histology to confirm the fiber position. The mice were deeply anesthetized by an overdose of ketamine/medetomidine, exsanguinated with phosphate buffered saline (PBS), perfused with 4% paraformaldehyde (PFA) in PBS. The brain was dissected from the skull and immersed in the 4% PFA for 12-24 hours at 4 °C. The brain was rinsed with PBS and sectioned (100 μm) by vibrating microtome (VT1000S, Leica).

Immunohistochemistry with TH antibody (AB152, Millipore Sigma; 1/750) was performed to identify dopamine neurons, with DsRed antibody (632496, Takara; 1/1000) to localize tdTomato expressing areas when tdTomato raw signal were not strong enough, and with GFP antibody (GFP-1010, Aves Labs; 1/3000) to localize GCaMP expressing areas when GCaMP raw signals were not strong enough. The sections were mounted on a slide-glass with a mounting medium containing 4’,6-diamidino-2-phenylindole (VECTASHIELD, Vector laboratories) and imaged with Axio Scan.Z1 (Zeiss).

### Behavior

After 5 days of recovery from surgery, mice were water-restricted in their cages. All conditioning tasks were controlled by a NIDAQ board and LabVIEW. Mice were handled for 2 days, acclimated to the experimental setup for 1-2 days including consumption of water from the tube, and head-fixed with random interval water for 1-3 days until mice show reliable water consumption. For odor-based classical conditioning, all mice were head-fixed, and the volume of water reward was constant for all reward trials (predicted or unpredicted) in all conditions (6 μl). Some condition contains mild air puff to an eye and the intensity of air puff was constant for all air puff trials (predicted or unpredicted; 2.5 psi). Each association trial began with an odor cue (last for 1 s) followed by 2 s delay, and then an outcome (either water, nothing, or air puff) was delivered. Odors were delivered using a custom olfactometer^42^. Each odor was dissolved in mineral oil at 1:10 dilution and 30 μl of diluted odor solution was applied to the syringe filter (2.7μm pore, 13mm; Whatman, 6823-1327). Different sets of odors (Ethyl butyrate, p-Cymene, Isoamyl acetate, Isobutyl propionate, 1-Butanol, 4-Methylanisole, (*S*)-(+)-Carvone, and 1-Hexanol) were selected for four groups: group 1 (4 animals for initial learning, 2 of which were used for reversal learning), group 2 (3 animal for initial learning, 1 of which was used for reversal learning), group 3 (2 animal for reversal learning (airpuff to reward) followed by repeated learning), and group 4 (3 animals for reversal learning (both nothing to reward and airpuff to reward) followed by repeated learning). Odorized air was further diluted with filtered air by 1:8 to produce a 900 ml/min total flow rate. A variable inter-trial interval (ITI) of flat hazard function (minimum 10s, mean 13s, truncated at 20s) was placed between trials. Each session was composed of multiple blocks (17-24 trial/block) and all trial types were pseudorandomized in each block. Each day, the mice did about 120-350 trials over the course of 25-75 min, with constant excitation from the laser and continuous recording.

Training for initial learning (Figure 1) started with an exposure to an odor chemical for the first time, using 4 types of trials; odor cue predicting 100% water, odor cue predicting 40% water/60% no outcome (nothing), odor cue predicting nothing (29.4% of all trials for each odor), and water without cue (free water) (11.8%) from day 1 to day 8, and odor cue predicting 80% water/20% nothing, odor cue predicting 40% water/60% nothing, odor cue predicting nothing (29.4% each), and free water (11.8%) on days 9 and 10. Neural activity was recorded in all ten sessions.

For reversal learning (Figure 2), mice were trained with classical conditioning for 12-24 days, and then, a nothing-predicting odor and/or airpuff-predicting odor were switched with an odor predicting high probability (80-100%) of water reward. More specifically, for reversal from both nothing- and puff-predicting odors to reward-predicting odors, 3 mice with GCaMP7f were first trained with odors A and B predicting 100% water, odor C predicting nothing, odor D predicting 100% airpuff (21.7% each), and free water (13.0%), from day 1 to day 12. On reversal day, odors A and B were switched with odors C and D, respectively. For reversal from nothing to reward with dopamine sensor-based recording, 3 mice used in the initial learning with DA sensor were trained for 9-13 more days (total 19-23 days), and then an 80% reward-predicting odor and a nothing-predicting odor were switched. For reversal of airpuff to reward, 2 mice with GCaMP6f were first trained with (day1-8) odor A predicting 100% water, odor B predicting 40% water, odor C predicting nothing, odor D predicting 100% airpuff (20.8% each), free airpuff (8.3%), and free water (8.3%) from day 1 to day 8, and then with odor A predicting 80% water, odor B predicting 40% water, odor C predicting nothing, odor D predicting 80% airpuff (20.8% each), free airpuff (8.3%), and free water (8.3%) from day 9 to day 23. On a reversal session, odor A was switched with odor D.

After 3 sessions of reversal learning, 3 mice with GCaMP7f were trained with 5 sessions of original conditioning (re-reversal; switching odor A and odor C, odor B and odor D again), before repeated learning (Figure 3). For repeated learning, an odor associated with 100% reward on re-reversal sessions (odorA) was replaced to a new odor, and the rest of trial types were kept exactly the same as re-reversal sessions. 2 mice with GCaMP6f were trained with repeated learning after 1 session of reversal learning. For repeated learning, an odor associated with high probability reward during a reversal session (odor C) was replaced to a new odor, and airpuff and free water trials were removed; new odor predicting 100% reward, odor B predicting 40% reward, and odor C predicting nothing (33.3% each).

## Data analysis

### Fiber-fluorometry

The noise from the power line in the voltage signal was cleaned by removing 58-62Hz signals through a band stop filter. Z-score was calculated from signals in an entire session smoothed with moving average of 50 ms. To average data using different sessions, signals were normalized as follows. The global change of signals within a session was corrected by linear fitting of signals and time and subtracting the fitted line from signals. The baseline activity for each trial (F0_each_) was calculated by averaging activity between -1 to 0 sec before a trial start (odor onset for odor trials and water onset for free water trials), and the average baseline activity for a session was calculated by averaging F0_each_ of all trials (F0_average_). dF/F was calculated as (F - F0_each_)/F0_average_. dF/F was then normalized by dividing by average responses to free water (0 to 1 sec from water onset). To detect activity peak, signals were first averaged over a sliding window of 3 trials. The activity peak during a cue/delay period was detected by finding a maximum response in moving windows of 20 ms that exceeds 2 × standard deviation of baseline activity (moving windows of 20 ms during -2 to 0 sec from an odor onset). The excitation onset during a delay period was detected by finding the first time point that exceeds 2 × standard deviation of baseline activity. To test the temporal shift of dopamine activity, we fitted the timing of activity peak or excitation onset to a trial number using a generalized linear model with exponential link function. The learning phase was defined as the duration until the timing of the activity peak shifts no more than 1ms/trial in the fitted exponential curve. To test the temporal shift of average activity of different animals, the first 40 trials in the first session were used. To detect activity peaks in the repeated learning, multiple local maximums of the activity with prominence of more than 2 × standard deviation of the baseline activity, more than 100 ms apart were detected. If the detected peaks were more than 2, the 2 peaks with the largest amplitude of activity were chosen. Then the last peak (or the detected peak if only one peak was detected) was used for the regression analysis to test for a temporal shift.

### Licking

Licking from a water spout was detected by a photoelectric sensor that produces a change in voltage when the light path is broken. The timing of each lick was detected at the peak of the voltage signal above a threshold. To plot the time course of licking patterns, the lick rate was calculated by a moving average of 300 ms window. The peak of anticipatory licking was detected at the maximum of lick rate during 0-3 s after odor onset. To detect anticipatory lick onset, the first and second licks were detected from 500 ms after odor onset because the onset of anticipatory licking is later than this period in well-trained animals. To binary score trials with or without anticipatory licking, trials with more than 4 licks during delay periods were defined as trials with anticipatory licking.

### Estimation of TD errors using simulations

To examine how the value and RPE may change within a trial and across trials, we employed a “complete serial compounds” approach to simulate animal’s learning^2^. We used eligibility traces to apply a TD(λ) learning method to learn state values (predicted reward value)^30^.

We considered two learning models, a TD(λ) model where 0< λ <1 and eligibility traces are used, and a model with Monte Carlo updates (or a TD(λ) model where λ =1). These models tiled a trial with 100 ms of serial different states. Each state from cue onset at time 0 to reward delivery at time 3 s has a weight to learn state value. Weights for all the states were initialized with 0. In each trial, eligibility traces for all the states were initialized with 0. At each time step, state value was calculated by

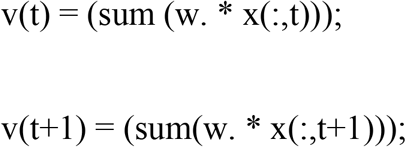

where v is a state value, w is a weight, x is a square matrix with a size of time steps, indicating states from cue to reward as 1, and otherwise 0. TD error was calculated by

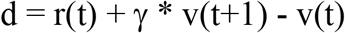

where d is TD error, r is reward, v is state value. Next, eligibility traces were updated by

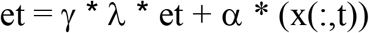

where et is eligibility traces for all the states, γ is a discounting factor from 0 to 1, λ is a constant to determine an updating rule and α is a learning rate. Then, weights are updated by

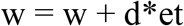

We used γ =0.98, λ =0.85, α=0.06, or λ =1, α =0.02 in Figure 4.

To mimic GCaMP signals recorded by fluorometry, obtained models were convolved with a filter of average GCaMP responses to water in dopamine axons^13^.

### Statistical analysis

All analyses were performed using custom software written in Matlab (MathWorks). All statistical tests were two-sided. A boxplot indicates 25th and 75th percentiles as the bottom and top edges, respectively. The center line indicates the median. The whiskers extend to the most extreme data that is not considered outlier. In other graphs, an error bar shows standard error. To test significance of the model fitting, *p*-value for the *F*-test on the model was calculated. One-sample t-test was performed to test the mean of a data set is not equal to zero. To compare difference of the mean between two groups, two-sample t-test was performed. *p*-value less than or equal to 0.05 was regarded as significant for all test.

## Materials, data and code availability

The fluorometry data will be shared at a public deposit source. The model code is attached as a source file. All other conventional codes used to obtain the result are available from the corresponding author. Original vectors, pAAV-TRE3G-WPRE and pAAV-TRE3G-GCaMP6f-WPRE will be deposit at Addgene. A mouse line Tg(Slc6a3-tTA)2Kftnk was deposit at RIKEN BioResource Center (BRC).

## Acknowledgements

We thank Iku Tsutsui-Kimura, Sara Matias, HyungGoo Kim, and Benedicte Babayan for technical assistance, Vanessa Roser and Sakura Ikeda for assistance in animal training, and Michael Bukwich and all lab members for discussion. We thank Catherine Dulac for sharing reagents and equipment. We thank Douglas Kim and GENIE Project, Janelia Farm Research Campus, Howard Hughes Medical Institute for pGP-CMV-GCaMP6f and pGP-AAV-CAG-FLEX-jGCaMP7f-WPRE plasmids, Edward Boyden, Media Lab, Massachusetts Institute of Technology for AAV5-CAG-FLEX-tdTomato and AAV5-CAG-tdTomato, Karl Deisseroth, Stanford University for pAAV-EF1a-DIO-hChR2(H134R)-EYFP-WPRE, and Yulong Li, State Key Laboratory of Membrane Biology, Peking University for AAV9-hSyn-DA2m. This work was supported by grants from National Institute of Health (U19 NS113201, NS 108740), and the Simons Collaboration on Global Brain (N.U.); Japan Society for the Promotion of Science, Japan Science and Technology Agency (R.A.); and Brain Mapping by Integrated Neurotechnologies for Disease Studies (Brain/MINDS) by AMED under grant number JP20dm0207069 (K.F.T).

## Author Contributions

RA and MWU designed experiments and analyzed data. RA collected data. RA and AY made constructs. KT made transgenic mice. The results were discussed and interpreted by RA, NU and MWU. RA, NU and MWU wrote the paper.

---

**Supplemental figure 1.**
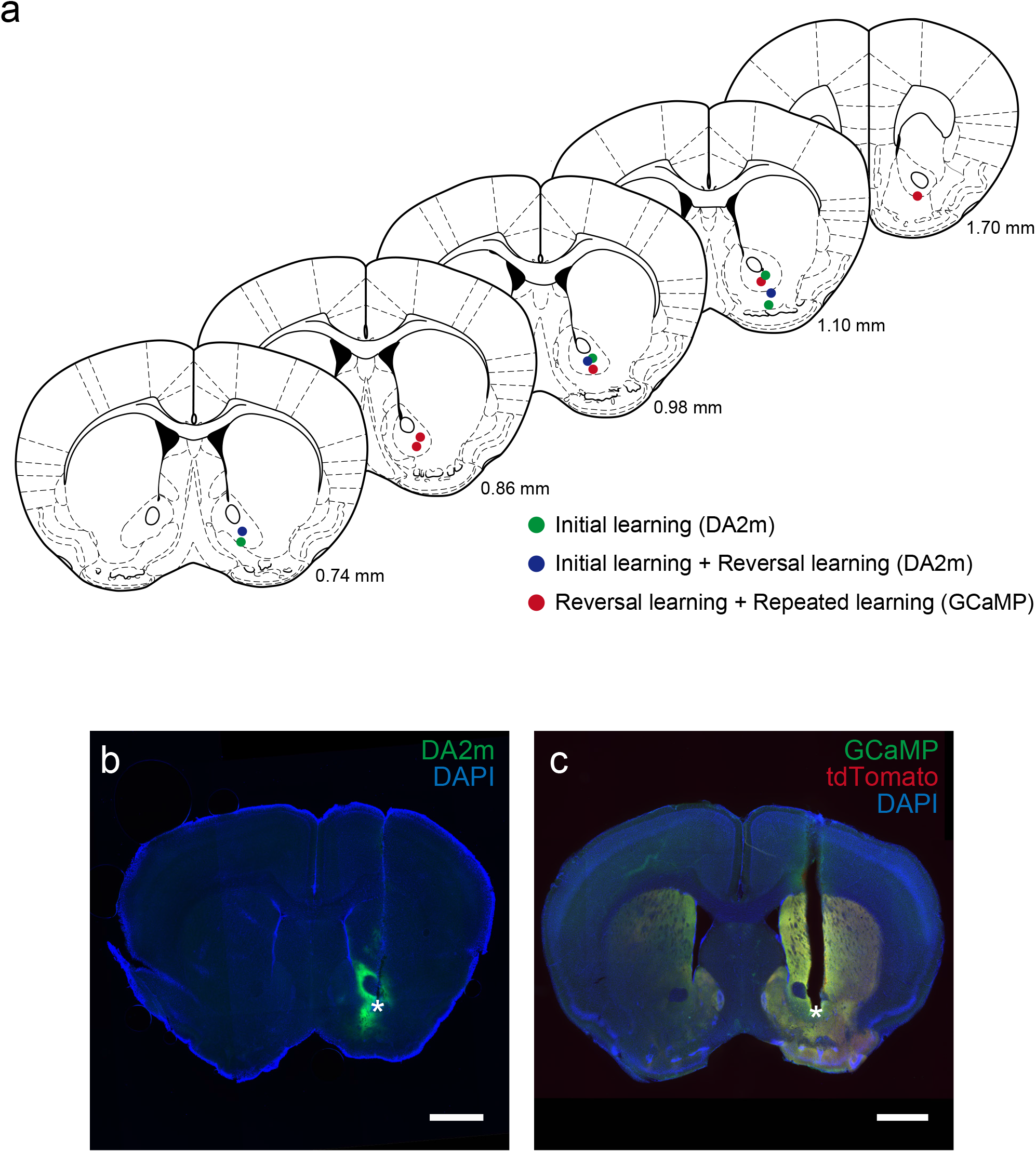
Recording sites for fiber-fluorometry. (**a**) Recording site for each animal is shown in coronal views (Paxinos and Franklin^42^). (**b**) Example coronal section of a recording site and DA2m (green) expression in VS. (**c**) Example coronal section of a recording site and GCaMP7f (green) and tdTomato (red) expression. Asterisks indicate fiber tip locations. Scale bars, 1 mm.

**Supplemental figure 2.**
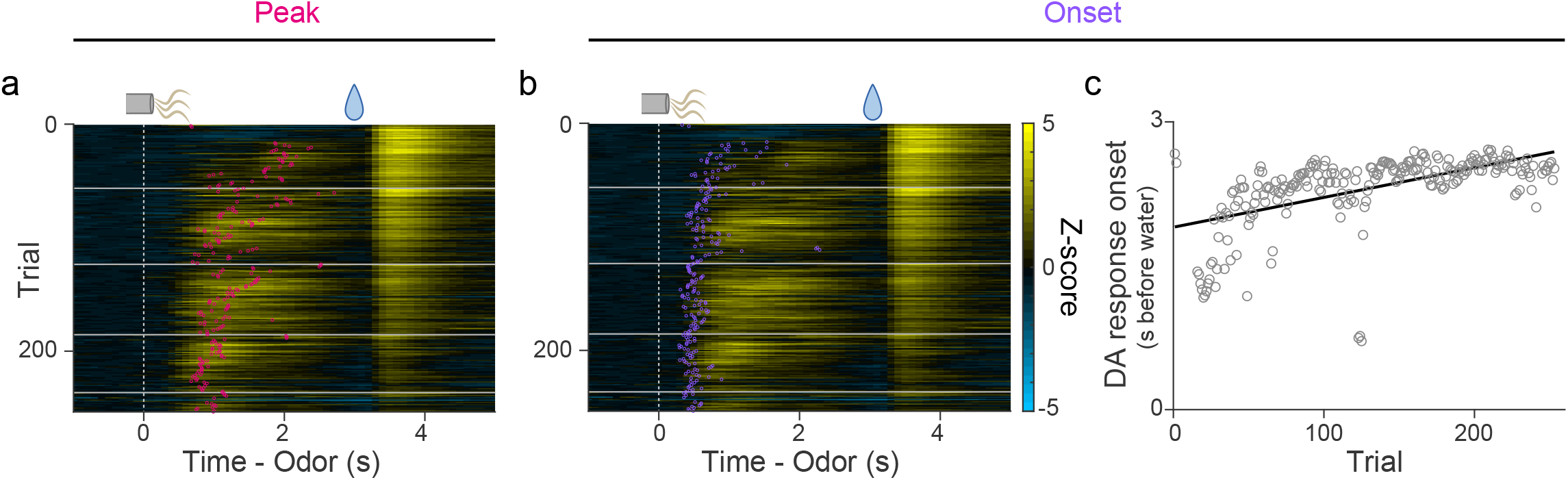
Peak and onset of dopamine responses to a reward-predicting odor in an example animal. (**a**) Dopamine sensor signal peaks during delay periods (red) overlaid on a heatmap of dopamine sensor signals in cued water trials. (**b**) Dopamine sensor response onset (purple) overlaid on a heatmap of dopamine sensor signals. (**c**) Linear regression of dopamine excitation onset with trial number.

**Supplemental figure 3.**
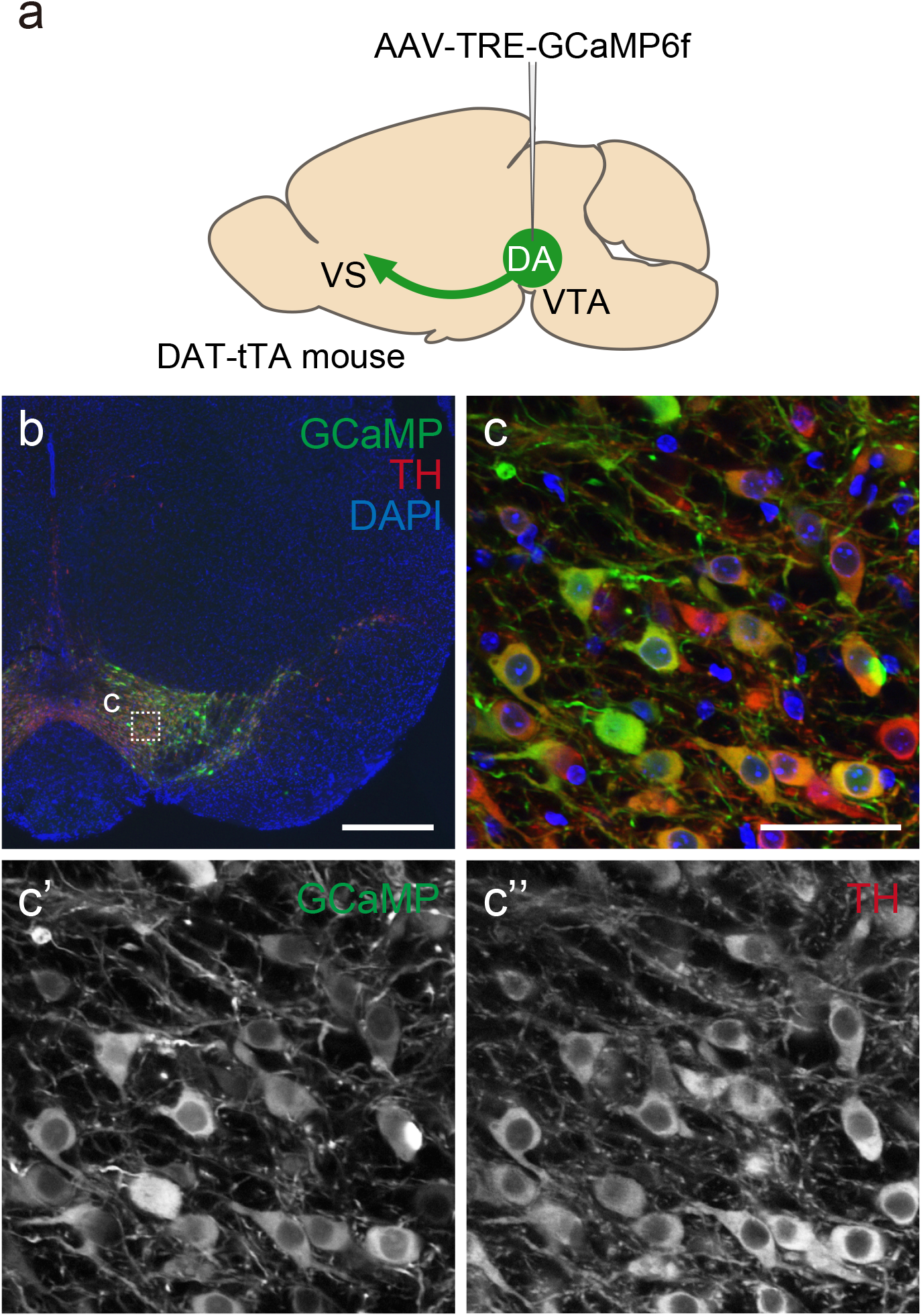
Dopamine neuron-specific GCaMP expression in DAT-tTA mice. (**a**) tTA-dependent AAV-GCaMP (AAV5-TRE3G-GCaMP6f) was injected into the VTA in 2 animals and used for reversal learning from airpuff to reward (Figure 2e-f) and repeated learning (Figure 3). (**b**) A coronal section of the midbrain in DAT-tTA mouse showing expression of GCaMP (green), and dopamine neurons labeled with antibody against tyrosine hydroxylase (TH) (red). The section was counterstained with DAPI (blue). Scale bar, 500 μm. (**c**) Magnified image of the patched area in VTA in (**b**), showing colocalization of GCaMP signals (green) and TH immunoreactivity (red). Single channel images for GCaMP and TH immunoreactivity are shown in (**c’**) and (**c’’**), respectively. Scale bar, 50 μm. Number of neurons positive for both GCaMP and TH immunoreactive signals of all GCaMP positive neurons is 98.7 ± 0.5% (mean ± sem, n = 3 animals, total 903 neurons) in VTA and 97.1 ± 0.8% (mean ± sem, n = 3 animals, total 229 neurons) in SNc.

**Supplemental figure 4.**
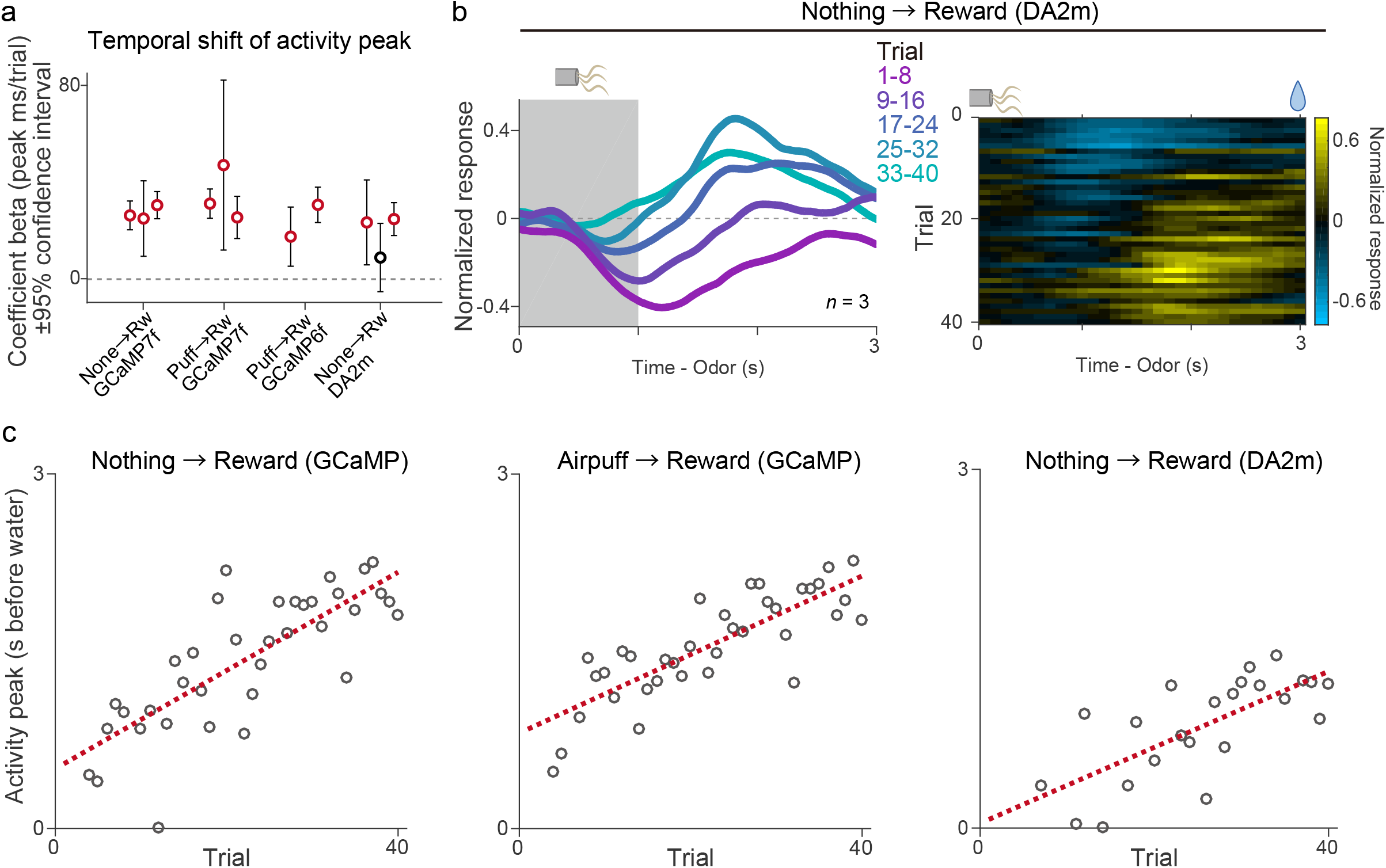
Temporal shift of activity during reversal learning. (**a**) The regression coefficients ±95% confidence intervals between activity peak timing and trial number in each animal under different experimental conditions. Red circles, significant (*p* ≤ 0.05) slopes. (**b**) Average dopamine activity (normalized to free water response) in response to a reward-predicting cue in the first session of reversal from nothing to reward (n = 3 animals with DA sensor). Each line shows 8 trials mean population neural activity across the session. Gray patch indicates the odor-presentation period. (**c**) Linear regression of peak timing of average activity with trial number during reversal from nothing to reward with GCaMP (left; n = 3 animals; coefficient beta = 41.6 ms/trial, *F* = 54, *p* = 1.5 × 10^−8^), from airpuff to reward with GCaMP (center; n = 5 animals; coefficient beta = 33.2 ms/trial, *F* = 66, *p* = 1.6 × 10^−9^), and from nothing to reward with dopamine sensor (right; n = 3 animals; coefficient beta = 31.8 ms/trial, *F* = 22, *p* = 1.3 × 10^−4^).

**Supplemental figure 5.**
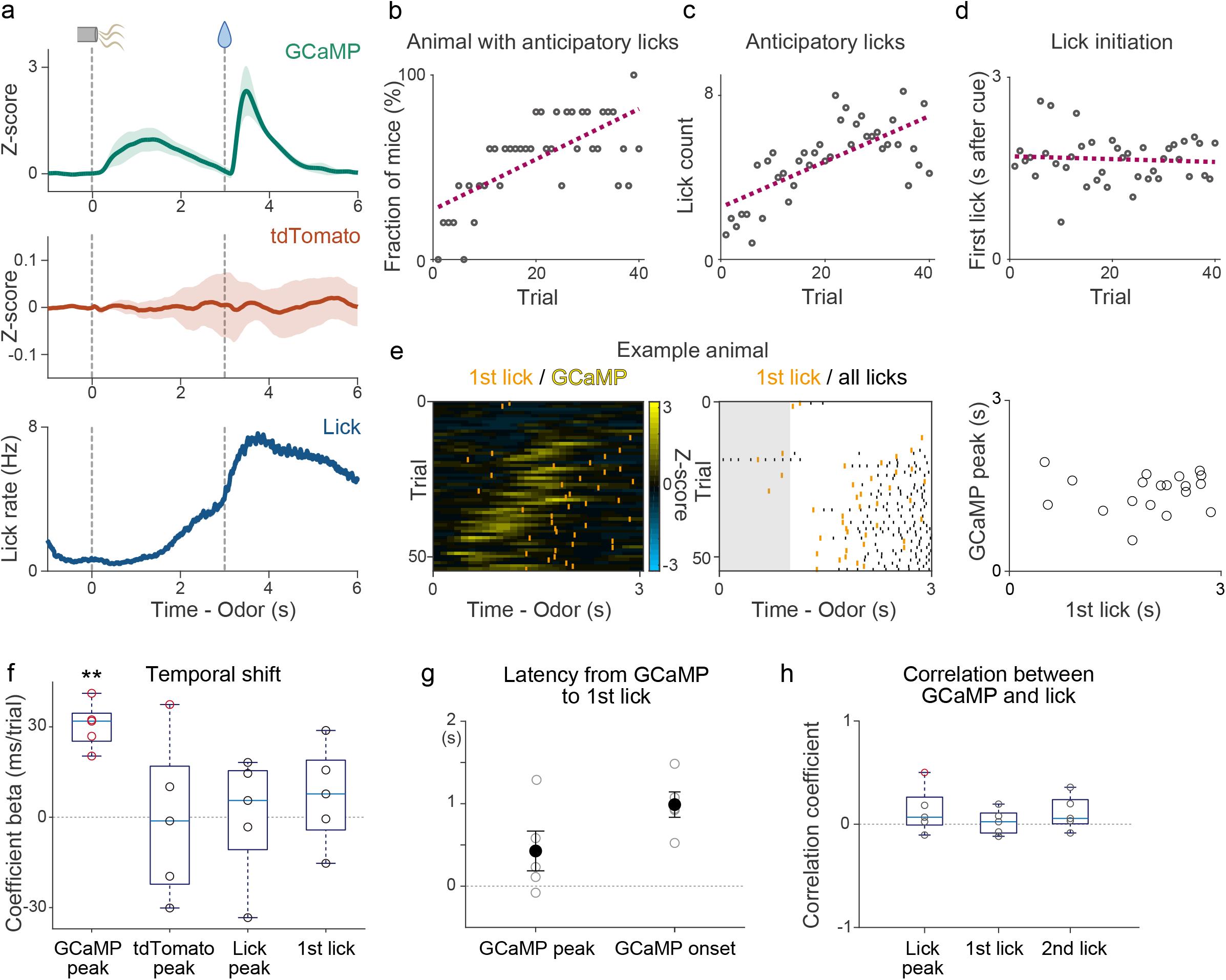
Comparison of dopamine axon GCaMP signal, control fluorescence signal, and licking. (**a**) GCaMP signals (top; green), tdTomato signals (middle; red), and lick counts (bottom; blue) recorded simultaneously in the first reversal session from airpuff to reward (mean ± sem). (**b**) Percentage of animals that show anticipatory licking during delay periods vs trial number. Regression coefficient beta = 1.37 (%/trial), *F* = 33, *p* = 1.2 × 10^−6^, *F*-test. (**c**) Average lick counts during delay periods vs trial number. Regression coefficient beta = 0.111 (lick/trial), *F* = 35, *p* = 7.8 × 10^−7^, *F*-test. (**d**) First lick timings vs trial number. Regression coefficient beta = -0.0023 (s/trial), *F* = 0.19, *p* = 0.66, *F*-test. (**e**) Relation between the first lick and GCaMP signals during the delay period in an example animal. Right, comparison between timing of GCaMP peak and timing of the first lick. (**f**) Linear regression coefficients for timing of GCaMP peak, tdTomato peak, lick peak and first lick with trial number (*t* = 8.9, *p* = 8.8 × 10^−4^ for GCaMP peak, *t* = -0.058, *p* = 0.96 for tdTomato peak, *t* = 0.038, *p* = 0.97 for lick peak, and *t* = 0.98, *p* = 0.38 for first lick; t-test). Red circles, significant (*p*-value ≤ 0.05, *F*-test). (**g**) Latency between GCaMP response and first lick (GCaMP peak to first lick; 427 ± 241 ms, and GCaMP response onset to first lick; 989 ± 154 ms, mean ± sem). (**h**) Correlation coefficients between timing of GCaMP response peak and lick peak (*t* = 1.3, *p* = 0.27, t-test) and lick onset (first lick, *t* = 0.53, *p* = 0.62, t-test, and second lick, *t* = 1.6, *p* = 0.19, t-test). Red circles, significant (*p*-value ≤ 0.05, *F*-test). n=5 animals.

**Supplemental figure 6.**
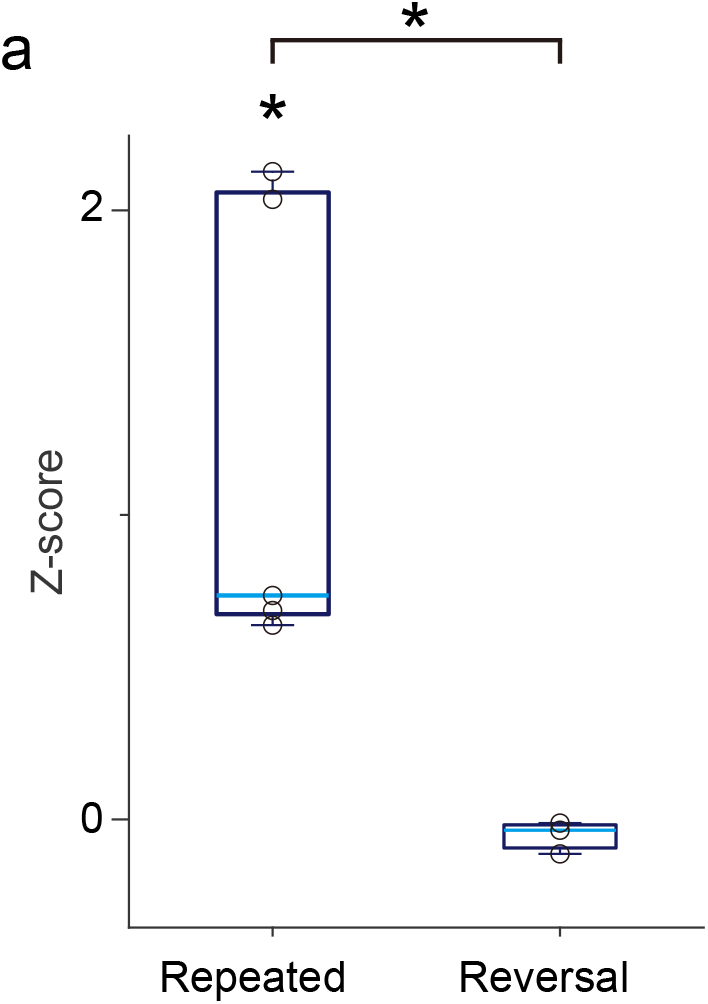
Dopamine axon response to odor in the first trial. (**a**) GCaMP response to a new odor (0-1 s after odor onset) in repeated learning (Figure 3) (n = 5 animals; *t* = 3.6, *p* = 0.022 compared to the baseline) and to an odor that had previously been associated with no outcome prior to reversal learning wherein it would become associated with reward (Figure 2) (n = 3 animals; *t* = -1.8, *p* = 0.22 compared to the baseline). Responses to odor in the first trial were significantly higher in repeated learning (*t* = 2.8, *p* = 0.030).

